# Beyond the known cuts: trypsin specificity in native proteins

**DOI:** 10.1101/2025.04.04.647035

**Authors:** Marcelo Gaspar, Bohdana Sokolova, Amirata Saei, José C. Marques, Roman A. Zubarev

**Affiliations:** Faculty of Exact Sciences and Engineering, University of Madeira, Campus Universitário da Penteada, 9020-105 Funchal, Portugal; ISOPlexis Centre for Sustainable Agriculture and Food Technology, University of Madeira, 9020-105 Funchal, Portugal; Division of Chemistry I, Department of Medical Biochemistry and Biophysics, Karolinska Institutet, SE-17 177 Stockholm, Sweden; Department of Microbiology, Tumor and Cell Biology, Karolinska Institutet, SE-17 177 Stockholm, Sweden; i3N-Institute for Nanostructures, Nanomodelling and Nanofabrication, Department of Physics, University of Aveiro, Campus Universitário de Santiago, 3810-193 Aveiro, Portugal

## Abstract

Trypsin is a serine protease that plays a pivotal role in protein digestion, being extensively used in various proteomics workflows due to its cleavage profile and specificity. Understanding trypsin’s enzymatic behavior is thus critical. We have employed Above-Filter Digestion Proteomics (AFDIP) to investigate trypsin’s cleavage preferences in HeLa cell lysates, preserving the proteins’ native state. We quantified over 18,000 unique peptides and correlated their emergence rates with cleavage window sequence motifs and physicochemical properties. Contrary to previous studies performed on denatured proteomes, we found that in native proteins cleavages at lysine residues were more abundant and faster than at arginine residues, and that physicochemical properties of the peptides affected their emergence times. These findings may need to be taken into account when interpreting the results of limited proteolysis experiments as well as when designing a food protein with extremely fast digestion times.

## INTRODUCTION

Trypsin is a serine protease important in both biology and research contexts (Kaur & Singh, 2022). Activated in the duodenum from its precursor trypsinogen by enzymatic removal of a 7-10 amino acid N-terminal peptide, trypsin is central to a variety of processes in human organism. Its most known function is to break down dietary proteins into smaller peptides and amino acids, which are further degraded by other proteases into single amino acids or very short peptides.

These peptides, typically not more than four amino acids in length, are efficiently absorbed by the intestinal mucosa (Fu et al., 2021). Additionally, trypsin amplifies protein digestion by activating other pancreatic zymogens, including chymotrypsinogen and proelastase, ensuring comprehensive breakdown of dietary proteins into absorbable forms. This enzymatic cascade is essential for nutrient assimilation and supports overall metabolic processes (Szabó et al., 2016; Whitcomb & Lowe, 2007). Trypsin is also involved in the kallikrein-kinin system, which helps regulate blood pressure (Vertiprakhov & Ovchinnikova, 2022). Besides that, trypsin activates protease-activated receptors (PARs), particularly PAR2, which play a role in inflammatory processes and immunological reactions (Amadesi & Bunnett, 2004). Through PAR activation, trypsin can influence the secretory function of the pancreas, stomach, and salivary glands (Kawabata et al., 2000; Namkung et al., 2004; Nishikawa et al., 2002). Trypsin participates in removal of dead skin cells and promote the growth of healthy tissue, aiding in wound healing (Xiang et al., 2024). Some evidence suggests trypsin may be involved in the pathogenesis of neurodegenerative diseases of the brain, though this requires further research (Hurley et al., 2015; Y. Wang et al., 2008).

In mass spectrometry (MS)-based proteomics, trypsin forms the basis for numerous analytical workflows. These rely on trypsin’s ability to produce peptides with optimal mass and charge properties for high-resolution MS, facilitating proteomics analysis (Kolsrud et al., 2012). By consistently generating well-defined peptide fragments, trypsin allows for accurate peptide mapping and comprehensive protein characterization, making it useful to uncovering protein structures and dynamics (Højrup, 2009). It also plays a role in elucidating complex biological processes, identifying biomarkers, and facilitating the discovery of novel therapeutic targets (e.g., Birhanu, 2023).

Trypsin exhibits remarkable specificity to the site of cleavages in proteins (Olsen et al., 2004). It is known to catalyze the hydrolysis of peptide bonds immediately following lysine (K) or arginine (R) residues, but not before proline (P) (so-called Keil rule K/R.P (Keil, 1992)). In MS analysis, this feature is useful as it guarantees the presence in a fully tryptic peptide of just one positively charged amino acid (Hunt et al., 1986; Olsen et al., 2004).

As the average occurrence of K and R together is ≈10%, the average size of a tryptic peptide is ≈10 amino acids, which corresponds to the molecular mass of 1.1-1.2 kDa. In electrospray ionization, such peptides would typically produce abundant 2+ and 3+ molecular ions, with one ionizing proton located on K or R, and the other protons – on a less basic sites, such as the N-terminal amine (Yang et al., 2012). These additional protons are easily mobilized in collisional activation, migrate along the backbone and initiate backbone cleavage, providing abundant sequence-specific information (Savitski et al., 2007).

Despite the widespread use of trypsin in proteomics, the subtleties of its specificity, especially when applied to complex protein mixtures, continues to demand further investigation. Although deep and efficient proteolysis is fundamental to the success of MS-based proteomics analysis (J. Wang et al., 2018), achieving it is far less trivial than appears at first glance. Optimal pH and temperature conditions are some of the parameters that are crucial for maximizing trypsin’s efficiency (Siepen et al., 2007). It is known that stable protein complexes (e.g., ribosome) and tight folding of the native protein structure can significantly affect the accessibility of cleavage sites, thus affecting the rate of digestion (Siepen et al., 2007). Autolysis of trypsin itself may reduce its effectiveness (Heissel et al., 2019), and therefore sequence-grade trypsin is heavily modified to reduce the self-proteolysis rate. In real-life proteomic experiments, despite the decades of protocol development and the use of modified trypsin optimized for specificity and stability, some polypeptide bonds amenable to trypsinolysis remain intact. This is why a typical proteomic data processing allows for up to two “missed cleavages” (Pirmoradian et al., 2013). Practically every proteomic dataset contains peptides with one or two sites uncleaved by trypsin, but the preferred motifs of these sites are not well established.

Instances of missed cleavages and incomplete protein digestion represent not only trypsin’s limitations, but also an opportunity for studying the complex interplay between protein conformation and post-translational modifications that seem to modulate the enzyme’s activity (Kim et al., 2014; Pan et al., 2014). For instance, the reduction of the hydrolysis rate at the site of drug binding can be used in chemical proteomics to identify the drug target (Holfeld et al., 2023). Recently, PELSA (peptide-centric local stability assay), a new proteolysis-based proteomics method for identifying protein targets and binding regions of diverse ligands, has been introduced (Li et al., 2025). PELSA employs a large amount of trypsin (enzyme-to-substrate ratio of 1:2 wt/wt) to generate peptides directly from treated/untreated lysates under native conditions. This approach allows for sensitive detection of ligand-induced protein local stability shifts on a proteome-wide scale. At the same time, the average degree of peptide bond cleavage in PELSA is quite low, which reflects in a smaller number of quantified peptides and proteins compared to full trypsinolysis. This also calls for better understanding of the trypsin digestion rate and specificity.

It is known that trypsin specificity can often be influenced by the neighboring residues near the cleavage site (Siepen et al., 2007). The accumulation of acidic residues adjacent to K or R sites can lead to diminished digestion rates (Korte et al., 2019; Šlechtová et al., 2015). The 2014 study by Pan et al. (Pan et al., 2014) provided a kinetic overview of trypsin-mediated protein digestion. They found that protein abundance and physicochemical properties, such as molecular weight (MW), grand average of hydropathicity (GRAVY), aliphatic index, and isoelectric point (pI) have no notable correlation with the priority of protein digestion. In that study Pan et al. used HeLa cell digest and analyzed three aliquots taken after 0.5, 2, and 18 h, representing fast, medium and slow digestion. The peptides in each aliquot were labeled by a corresponding isotopically different dimethyl label, multiplexed and fractionated into 6 fractions. Each fraction was analyzed by LC-MS/MS with high-resolution MS and low-resolution MS/MS. This resulted in the quantification of 10,483 unique peptides from 2,270 proteins. Almost 43 % of peptides had at least one missed cleavage site. Based on the abundances of the isotopically labeled molecular ions, all peptides were classified as either early generated or late generated. The sites of cleavage were classified into four groups: as very fast, fast, slow, and very slow sites. It was confirmed that K/R cleavage sites surrounded by neutral amino acid residues are quickly cut, while those with neighboring charged residues (D/E/K/R) or P are hydrolyzed slowly. Another finding was that trypsin cleaves the C-terminal to R at a with higher rate than for K.

It should be noted that in the work of Pan et al., proteins were denatured in 8 M urea and S-S bonds were reduced before digestion, as customary in shotgun proteomics. This, however, could have affected the results and explained why that team found no correlation with physicochemical properties of proteins. Protein denaturation before digestion is not an obligatory feature – for instance, in limited proteolysis (Holfeld et al., 2023) and other proteomic approaches that aim at probing protein structure, protein-protein and protein-drug interaction (Bennett et al., 2022; Yu & Huang, 2018), denaturation is avoided. For the purpose of understanding better how proteins are digested in guts of animals and humans, denaturation before digestion is also undesirable.

Having the latter aim, we set to investigate which specific sequence motifs challenge trypsin’s efficacy and which, to the contrary, provided fastest cleavage in proteins retaining their native conformation. While Pan et al. reported overall cleavage trends for trypsin, their study of denatured samples did not take into account possible conformational effects observed under native conditions. By preserving the natural protein structures, the current study provided a more physiologically relevant perspective on trypsin’s enzymatic activity.

Here we performed Above-Filter Digestion Proteomics (AFDIP) (Sokolova et al., 2025) on the proteome extracted from HeLa cells (Fig. 1). With a 1 h interval, the products of trypsinolysis are filtered through a membrane filter with a 3 kDa molecular weight cut-off (MWCO). After that the filtrate is collected and additional buffer is added to the undigested proteins above the filter. After 8 h of digestion, all collected filtrates with released peptides are labelled with the isobaric tandem mass tag (TMT). Such procedure is performed for at least 3 replicates. The labeled filtrate fractions are then combined, S-S bonds are reduced, and the multiplexed samples are separated into 24 fractions using off-line 2D chromatography for deeper analysis. Each fraction is then analyzed by LC-MS/MS with high resolution both in MS and MS/MS, for reliable peptide sequence identification. The center of gravity (CoG) of the elution curve for each peptide is calculated as:

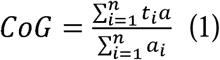

where *t* is a timepoint (1–8 h), and *a* is the relative abundance of a peptide of interest. The CoG values are then correlated with the peptide sequence and serve as an indicator of average digestion time, with smaller CoG values representing faster cleavage rates. By aligning tryptic peptide sequences relative to cleavage sites and extending ±6 residues on either side, we could assess sequence motifs while simultaneously exploring correlations with peptide physicochemical properties such as molecular weight, GRAVY, aliphatic index, and pI. This integrated approach provided a comprehensive understanding of trends influencing digestion efficiency. It also serves as a stepping stone for further exploration of trypsin activity under varying conditions, providing valuable insights for the optimization of proteomics workflows.

**Fig. 1.**
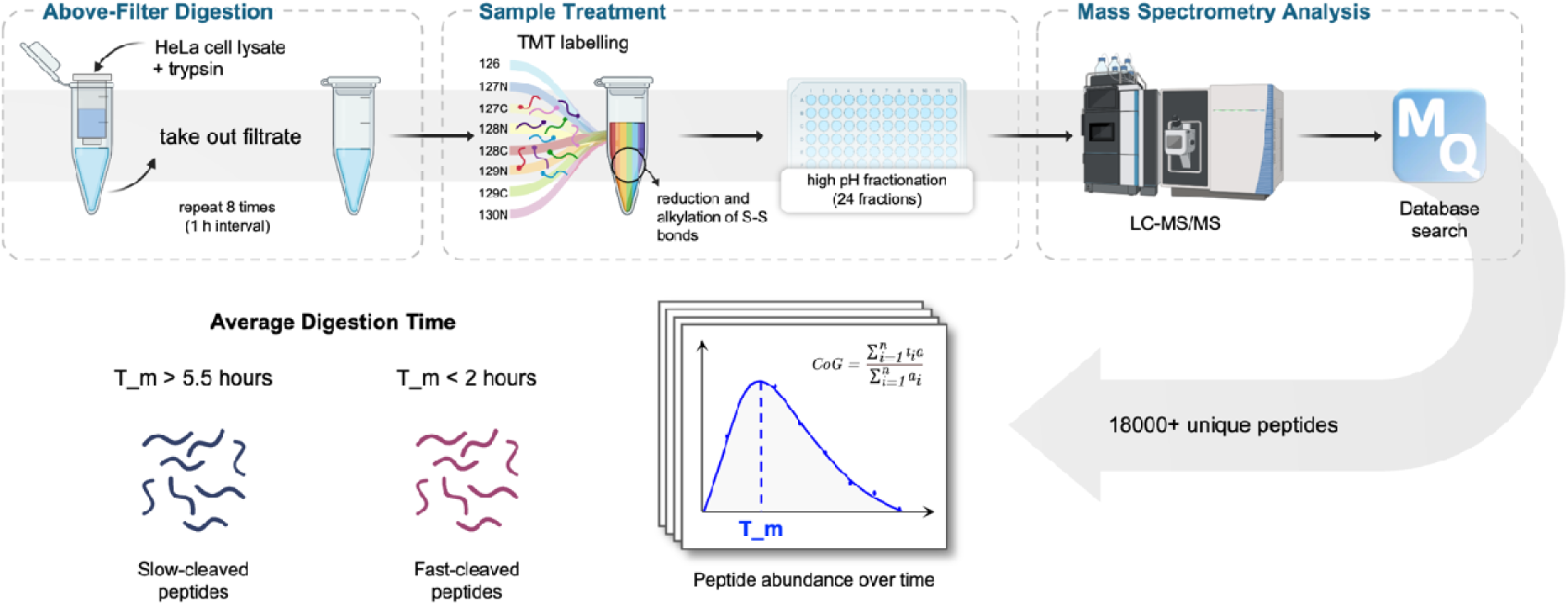
Workflow for the Above-Filter Digestion Proteomics (AFDIP) method. (Sokolova et al., 2025). Limited proteolysis is performed with trypsin-mediated digestion of HeLa cell lysates above a low-mass (3 kDa) molecular weight cut-off (MWCO) filter. At pre-set equidistant time points following the beginning of digestion, the digest solution is centrifuged for fixed time. The filtrate containing the peptides is collected, while the same amount of new buffer with trypsin is added to the above-filter solution containing under-digested proteins and the remaining protease. This procedure is performed 8 times with 1 h intervals during 8 h until digestion is completed. The collected filtrates from each hour are labeled with isobaric tandem mass tags (TMT16) to facilitate downstream multiplexed analysis. After labeling, disulfide bonds are reduced and alkylated. Peptides are fractionated with each fraction subsequently analyzed by liquid chromatography-tandem mass spectrometry (LC-MS/MS), with peptide identification performed using database search. Each identified peptide is classified based on its average digestion time (T_m), calculated as the center of gravity (CoG) of their elution curves over time. Slowly cleaved peptides (T_m > 5.5 h) and quickly cleaved peptides (T_m < 2.0 h) are analyzed separately to examine digestion dynamics and identify sequence motifs and physicochemical correlations. This comprehensive approach yields over 18,000 unique peptides, providing insights into trypsin specificity and peptide emergence trends. Figure created with BioRender (https://BioRender.com)

## RESULTS

In total, 18616 unique peptides were identified belonging to 3087 unique proteins. Of these peptides, 9402 ended with lysine (K) and 8724 ended with arginine (R). There were 13982 fully tryptic peptides (∼75.1%) and 4634 semi-tryptic peptides (∼24.9%). Of the latter, 4192 molecules (∼22.5%) were with one and 442 (∼2.4%) with two missed cleavages. The CoG distribution of these proteins is shown in Fig. 2A. Most peptides cluster around a digestion time of 3–5 h, with extremes representing fast-cleaved and slow-cleaved peptides. Notably, the distribution is somewhat asymmetric, with a steeper slope on longer digestion times. This could be due to a two-phase protein degradation process, with the native structure degrading in the first phase and denatured proteins being cleaved in the second, faster phase. The presence of two modes did not however affect significantly the results of our study.

**Fig. 2.**
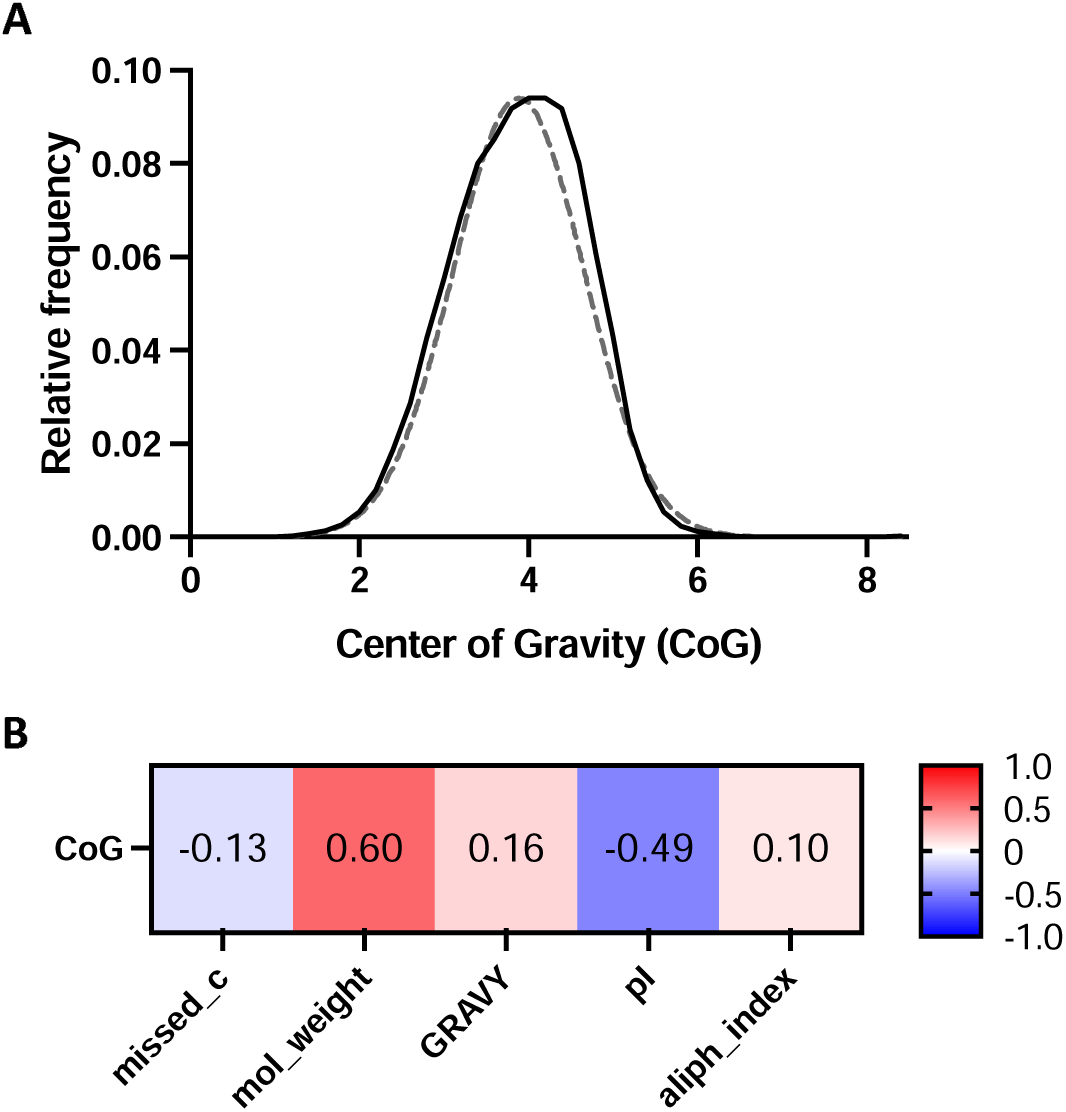
Peptide distribution and physicochemical correlations based on the calculated Center of Gravity (CoG) values. CoG is used as a metric for the average digestion time (in hours) of each quantified peptide across the digestion time course. **(A)** shows the density distribution of 18616 peptides as a function of their calculated CoG values (binned in 0.2 h intervals) – the dashed line represents a theoretical symmetric normal distribution centered at the observed mean CoG (3.9 h) and scaled to the maximum observed relative frequency. **(B)** shows Pearson correlation values of the peptides’ CoG with the peptide’s number of missed cleavages (missed_c) and other features such as: molecular weight (mol_weight), grand average of hydropathy (GRAVY), isoelectric point (pI) and aliphatic index (aliph_index). All correlations shown are statistically significant (p < 0.0001).

CoGs of peptides showed significant positive correlation with peptide mass (Fig. 2B). This could be because the larger the tryptic peptide, the more likely it is to accommodate negatively charged residues, which repel trypsin. On the other hand, isoelectric point (pI) anti-correlated with CoG, which is also understandable: high pI means the abundance of positively charged residues that attract trypsin, and hence faster cleavage. Other peptide features, such as the grand average of hydropathy (GRAVY) and aliphatic indexes, showed much weaker correlation with CoG. Somewhat surprisingly, even the presence or absence of missed cleavages did not correlate significantly with CoG.

### Preferred and avoided motifs

Motif analysis revealed distinct preferences and avoidances in amino acid sequences surrounding the tryptic cleavage sites (Fig. 3). Peptides were categorized into two digestion-speed groups: fast-emerging peptides (T_m < 2.0 h, including 252 cleavage sequences associated with 131 peptides) and slow-emerging peptides (T_m > 5.5 h, comprising 354 cleavage sequences associated with 185 peptides).

**Fig. 3.**
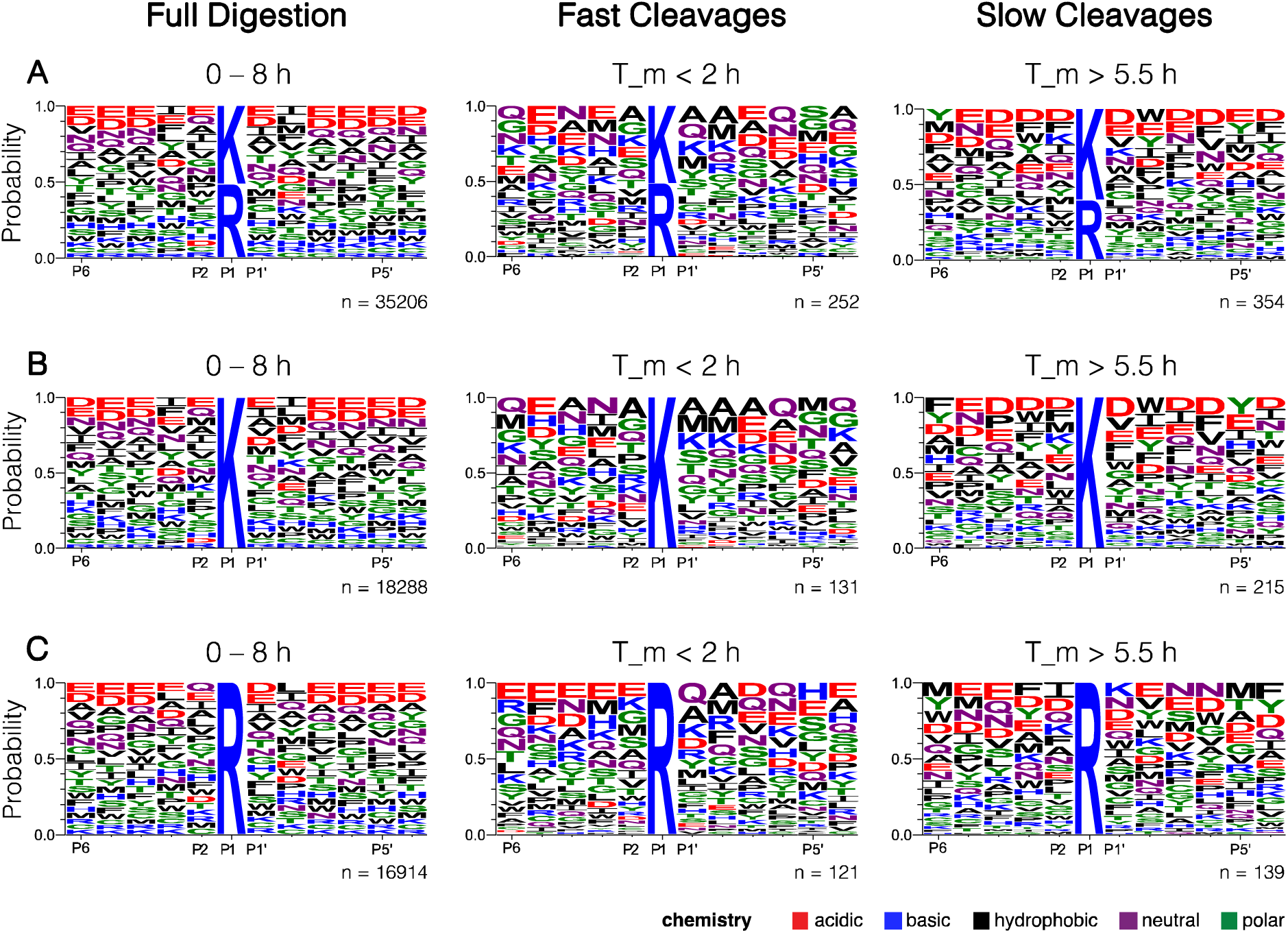
Sequence logos representing the frequency distribution of the amino acid residues surrounding the cleavage sites. Logos are centered at the cleavage site, which occurs between P1 and P1’, for **(A)** full sequence windows, **(B)** windows considering K at P1, and **(C)** windows considering R at P1.

As expected, the sequence logos generated from the corresponding normalized cleavage windows (±6 residues around the cleavage site) show a clear enrichment of K and R residues at the P1 position, consistent with trypsin’s substrate specificity. In some earlier studies, trypsin has been observed to cleave at R residues with greater efficiency than at K (Pan et al., 2014).

However, in our case, about 52% of the cleavages (18288 of the total 35206) occurred after K. Interestingly, this frequency increased to ∼61% (215 out of 354) in the slower cleaved sequences. In fast-emerging peptides, cleavages after K were slightly more frequent than R (131 out of 252, ∼52%). Adjacent to the cleavage site, hydrophobic residues (alanine at P2 and P1’, for instance) and other non-acidic residues were more frequent in fast-emerging sequences. At the same time, acidic residues (e.g., aspartic acid (D) and glutamic acid (E)) seem to be avoided there, showing instead a higher prevalence in slower cleaved sequences. These observations align with the established findings on trypsin cleavage efficiency, which show a particular reduced cleavage efficiency when K and R residues are surrounded by negatively charged amino acids, likely due to unfavorable interactions with the enzyme’s active site (Korte et al., 2019; Mori et al., 2022; Nickerson & Doucette, 2022; Ye et al., 2014).

### Correlation with MW

The noticed on Fig. 2B tendency for peptides with higher molecular weights to emerge later (positive correlation with CoG) needed further investigation. A more detailed picture is presented in Fig. 4. Here, the fast-emerging and slow-emerging groups of peptides only partially overlap, with the majorities of the distributions clearly separated. Note that Pan et al. (2014) did not find such a tendency. This could be explained by the fact that in their study, proteins were denatured, which apparently resulted in uniform accessibility of trypsin to potential cleavage sites. Under native conditions, as in our study, peptide size seems to significantly influence the peptide emergence dynamics. Besides the already mentioned possibility of negative charge influence, larger peptides exhibit lower mobility and enhanced tendency of interacting with other polypeptides, which may also delay their emergence.

**Fig. 4.**
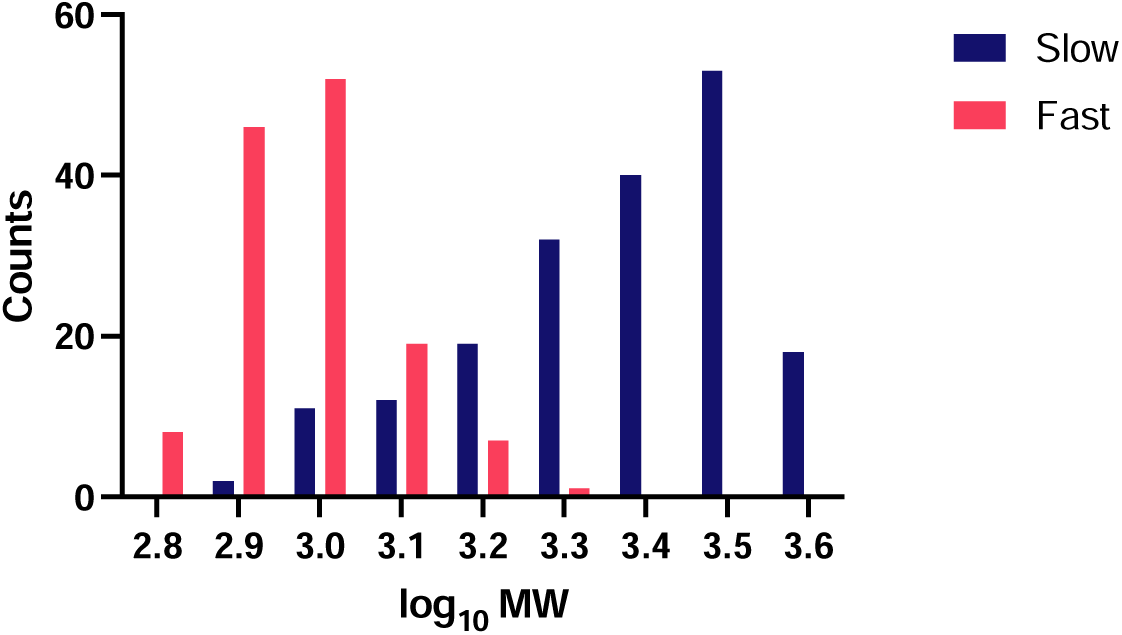
**Log_10_ molecular weight (MW) distribution of early emerging (fast cleaved) and late emerging (slow cleaved) peptides.**

### Correlation with GRAVY

The relationship between CoG and the GRAVY score showed a slight positive correlation (Fig. 2B), suggesting that peptides with higher GRAVY values (more hydrophobic) tend to emerge somewhat later. As shown in Fig. 5, the distribution of GRAVY values for early and late-emerging peptides indeed suggests some differentiation, with faster-cleaving peptides displaying a tendency toward more hydrophilic values (more negative GRAVY scores; average value −1.2), while slower-cleaving peptides tend to be less hydrophilic (average value −0.5). This suggests that hydrophilic cleavage sites, often surface-exposed in native protein structures, are more accessible to trypsin during digestion. Conversely, hydrophobic regions are more likely to be buried inside the protein core, requiring structural rearrangements or partial unfolding to expose these sites to trypsin. These findings also diverge from the observations under denatured conditions (Pan et al., 2014), where hydrophobicity shows minimal influence.

**Fig. 5.**
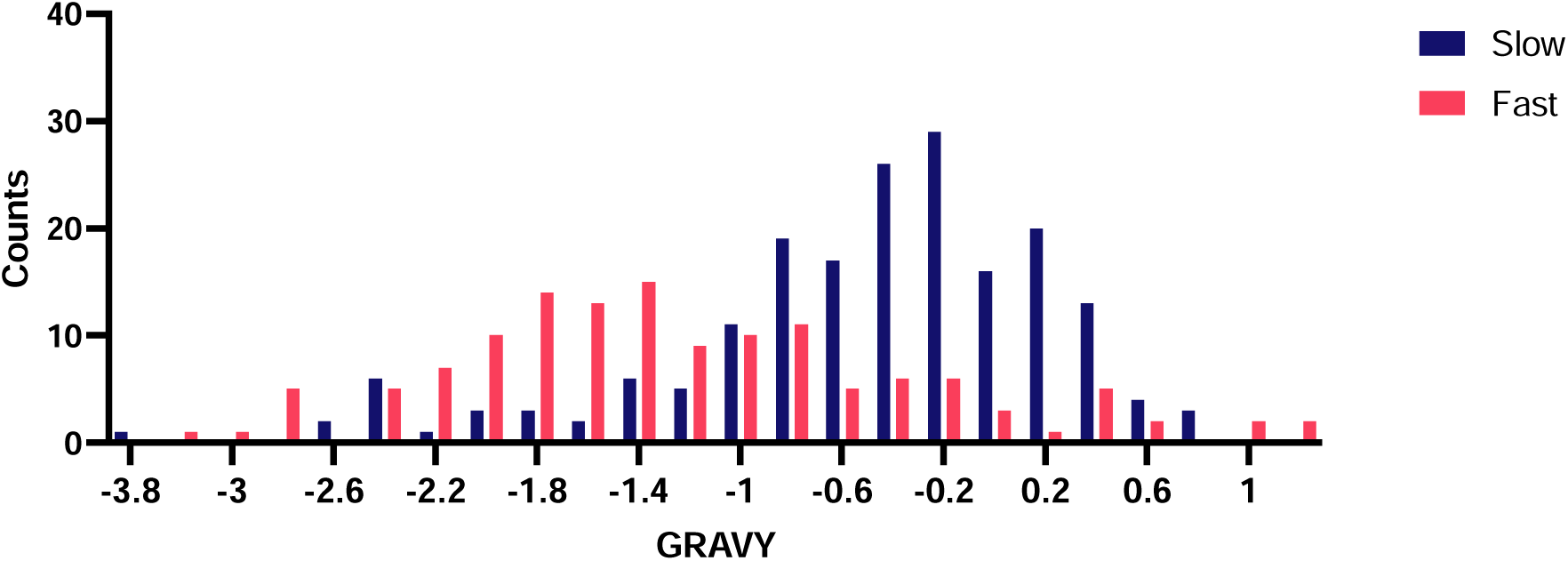
**Distribution of grand average of hydropathy (GRAVY) of early emerging (fast cleaved) and late emerging (slow cleaved) peptides.**

### Correlation with aliphatic index

Similarly, data shown in Fig. 6 seem to highlight a slight preference of trypsin to release aliphatic peptides, further supporting the enzyme’s bias toward certain structural and compositional attributes of proteins. Once again, the study with denatured proteins (Pan et al., 2014) could not capture this aspect.

**Fig. 6.**
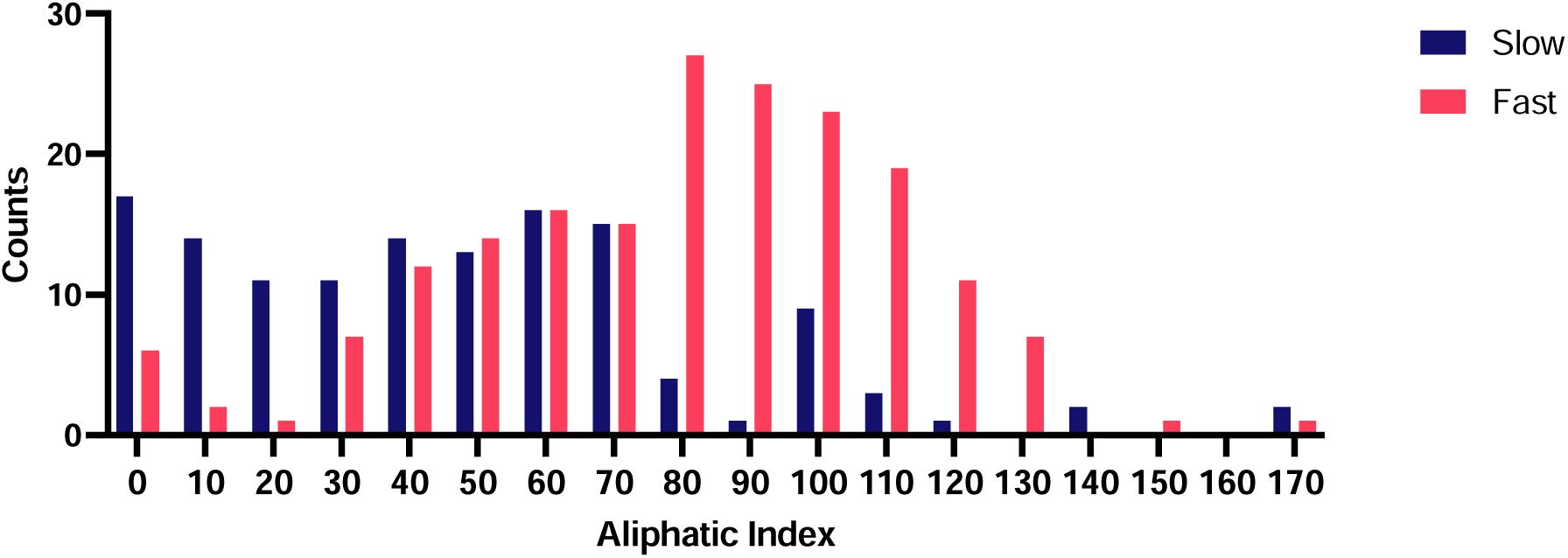
**Distribution of the aliphatic index values of early emerging (fast cleaved) and late emerging (slow cleaved) peptides.**

### Correlation with pI

The analysis of peptide counts varying with isoelectric point (pI) (Fig. 7) revealed that peptides with a pI around 4 were highly abundant. Absolute majority of these peptides emerge later, suggesting that their acidic nature delays cleavage. Conversely, peptides with pI values greater than 6 tend to emerge earlier. Pan et al. found no notable correlation of pI with digestion priority of proteins (Pan et al., 2014).

**Fig. 7.**
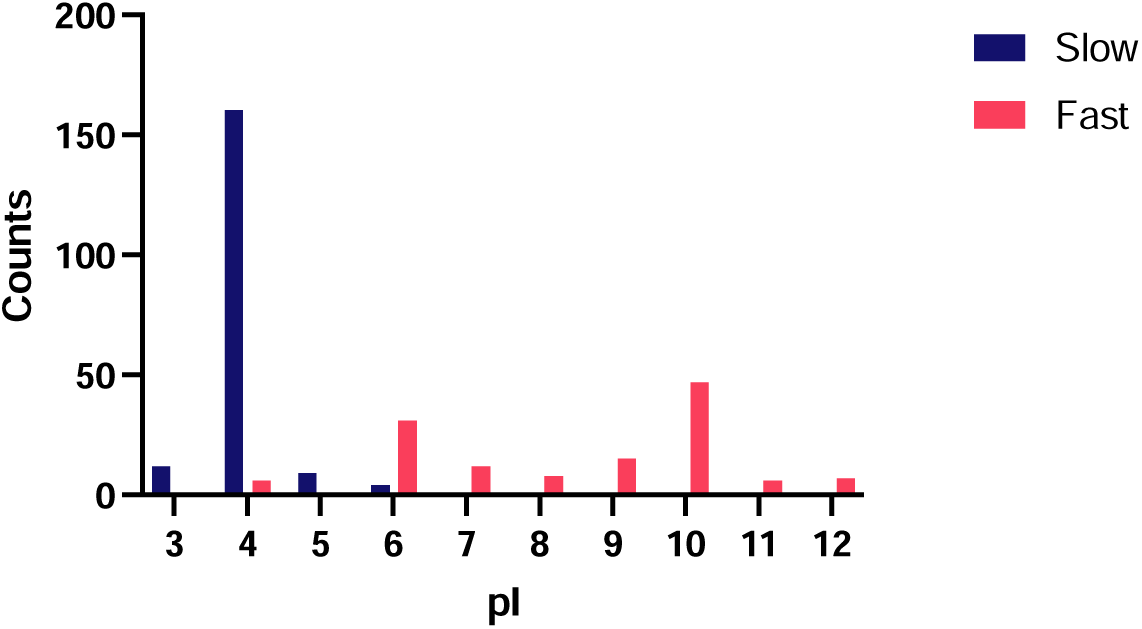
**Distribution of the theoretical isoelectric point (pI) values for early emerging (fast cleaved) and late emerging (slow cleaved) peptides.**

## CONCLUSIONS

Trypsin’s primary specificity (K/R) appears to be modulated by neighboring amino acids other than proline at P1’. We show that charged residues near the scissile bond slow trypsin hydrolysis and can reduce cleavage efficiency, whereas more “friendlier” neighboring amino acids (small or neutral residues) appear to promote a faster hydrolysis. Beyond the immediate sequence motif, the broader physicochemical properties of peptides and proteins can influence how quickly trypsin cleaves at a given site – especially under limited proteolysis conditions. Although in a denatured proteome such intrinsic properties seem to play a minimal role, our data collectively demonstrate that MW, GRAVY, aliphatic index, and pI, appear to influence digestion dynamics under native protein conditions. These factors may need to be taken into account when interpreting the results of limited proteolysis experiments. Additionally, these factors identified as influencing cleavage rates under limited proteolysis conditions may directly translate to how proteins digest in the gut and could help guide the design of food protein with extremely fast digestion times.

## METHODS

### Above-Filter Digestion procedure

The experimental design followed a limited proteolysis approach above a low-mass (3 kDa) cut-off filter – Above-Filter Digestion Proteomics (AFDIP, Fig. 1) (Sokolova et al., 2025). Briefly, at pre-set equidistant time points following the beginning of digestion, the digest solution of HeLa cell lysate was centrifuged for a fixed time. The filtrate, containing digested peptides, was then collected and an equal volume of fresh buffer was added to the above-filter solution containing under-digested proteins and the protease. This procedure was performed 8 times with 1 h intervals during 8 h. The collected filtrates were labeled with tandem mass tags (TMT16) and pooled together. Then, reduction and alkylation of disulfide bonds and desalting of the samples was performed. Samples were then fractionated to reduce sample complexity prior to analysis using liquid chromatography-tandem mass spectrometry (LC-MS/MS). For every identified peptide, an elution curve was constructed based on peptide abundance and digestion time. The center of gravity (CoG) of the curve was calculated to represent the average digestion time (T_m). For full details on proteome extraction, sample preparation, LC-MS/MS analysis and data processing, please refer to Sokolova et al. (2025).

### Cleavage motif analysis

Similar to Pan et al. (2014), raw sequences were first centered at the cleavage site and extended by 12 residues (±6 residues to both ends) to form cleavage windows with the format [P6’ P5’ P4’ P3’ P2’ P1’ P1 P2 P3 P4 P5 P6]. Sequences that could not be extended on either the N- or C-terminal sides were excluded from the analysis. To account for the natural abundance of amino acids and eliminate bias in sequence analysis, the raw frequencies were also normalized.

Specifically, amino acid frequencies at each position were multiplied by 1000 and divided by their natural occurrence in the human proteome (Proteome ID: UP000005640) as provided by UniProt. This normalization process ensured that overrepresented amino acids were adjusted appropriately, providing a corrected representation of positional preferences. Data was first processed in Microsoft Excel and then with RStudio (version 2024.04.2+764), using a custom script built with the Biostrings R package version 2.72.1 (Pagès et al., 2024). The script organizes raw FASTA files and computes positional amino acid frequencies, generating a file that accurately reflected the normalized distribution. These frequencies were subsequently converted into pseudo-sequences, representing the normalized data, solely for the purpose of generating the final sequence logos. These were created using WebLogo’s sequence logo generator (version 3.7.12) (Crooks et al., 2004).

### Physicochemical properties and correlation analysis

Additional physicochemical properties of peptides were computed using the R package Peptides version 2.4.6 (Osorio et al., 2015). Specifically, grand average of hydropathicity (GRAVY: hydrophobicity using the Kyte-Doolittle scale), isoelectric point (pI: pI), aliphatic index (aIndex) and molecular weight (mw) were calculated for each sequence. Pearson correlation analyses between these properties and peptide CoG values were performed in GraphPad Prism version 10.2.1 (GraphPad Software, San Diego, CA, USA).

